# Clostridium difficile Toxin B activates the NLRP3 inflammasome in human macrophages, demonstrating a novel regulatory mechanism for the Pyrin inflammasome

**DOI:** 10.1101/2021.07.07.451422

**Authors:** Matthew S.J. Mangan, Friederike Gorki, Karoline Krause, Alexander Heinz, Thomas Ebert, Dieter Jahn, Karsten Hiller, Veit Hornung, Marcus Mauer, Florian I. Schmidt, Ralf Gerhard, Eicke Latz

## Abstract

Pyrin is a cytosolic immune sensor that forms an inflammasome when bacterial virulence factors inhibit RhoA, triggering the release of inflammatory cytokines, including IL-1β. Gain of function mutations in the MEFV gene encoding Pyrin cause auto-inflammatory disorders, such as familial Mediterranean fever (FMF) and Pyrin associated auto-inflammation with Neutrophilic Dermatosis (PAAND). To precisely define the role of Pyrin in detecting pathogen virulence factors in relevant human immune cells, we investigated how the Pyrin inflammasome response was initiated and regulated in monocyte-derived macrophages (hMDM) compared to human monocytes. Unlike monocytes and murine macrophages, we determined that hMDM failed to activate Pyrin in response to known Pyrin activators *Clostridioides difficile* (*C. difficile*) toxins A or B (TcdA or TcdB). In contrast, TcdB activated the NLRP3 inflammasome in hMDM. Notably, we ascertained that the Pyrin inflammasome response could be re-enabled in hMDM by prolonged priming with either LPS or type I or II interferons, and required an increase in Pyrin expression. These data demonstrate an unexpected redundancy in detecting these toxins by inflammasome sensors.

## Introduction

Inflammasome-forming proteins are cytosolic sensors that mediate a post-translational inflammatory response to pathogens or cell stress. Upon detecting a stimulus, these sensors oligomerize and recruit the adapter protein ASC, enabling activation of caspase-1. Active caspase-1 then mediates cleavage and release of pro-inflammatory cytokines including IL-1β, and pro-inflammatory cell death by cleaving the pore-forming molecule gasdermin D [1]. Notably, rather than relying solely on direct detection of pathogens, some inflammasome sensors have instead evolved to detect infection or cellular stress by monitoring disruption of cellular homeostasis [2]. A leading example of this is the Pyrin inflammasome, encoded by the *MEFV* gene, which is activated in response to inhibition of RhoA [3]. This small G protein controls cytoskeletal rearrangement and is essential for immune cell migration and phagocytosis, amongst other functions [4]. Unsurprisingly, given its role in fundamental cellular processes, RhoA is a target of numerous bacterial virulence factors from pathogenic bacteria, including *Clostridium difficile* toxins A and B (TcdA and TcdB, respectively) [5]. Another prominent sensor of homeostasis is the NLRP3 inflammasome, which detects a wide range of events that converge to cause either mitochondrial dysfunction or loss of osmotic control of the cytosol [6].

Due to their high inflammatory potential, inflammasome forming sensors are strictly regulated by post-translational controls. Under resting conditions, Pyrin is maintained in an inactivate state by two distinct mechanisms regulated by RhoA signaling. The most well-characterized of these is phosphorylation of Pyrin at residues Ser208 and Ser242 by PKN1/2, members of the PKC superfamily [7–10]. Phosphorylation of Pyrin enables the subsequent binding of 14-3-3 proteins, which sequesters Pyrin in an inactive state. The second mechanism regulating Pyrin activation is less well understood, but is defined by a requirement for microtubule dynamics [9]. Pretreatment with colchicine, a microtubule destabilization agent, inhibits Pyrin activation but does not prevent dephosphorylation of Pyrin, demonstrating that these two regulatory mechanisms are mutually exclusive. Interestingly, human and mouse Pyrin share these regulatory mechanisms, although murine Pyrin does not contain the C-terminal B30.2 domain due to a frameshift mutation [11].

Understanding the regulatory mechanisms governing Pyrin is particularly important as mutations in the *MEFV* gene cause the hereditary auto-inflammatory disorder, Familial Mediterranean Fever (FMF). FMF is characterized by recurrent attacks of fever, serositis, abdominal pain and can, over time, cause secondary AA amyloidosis, leading to kidney failure [12]. Mutations linked to FMF are mostly amino acid substitutions in the C-terminal B30.2 domain [13]. Though the function of the B30.2 domain is still relatively unclear, these mutations are suggested to perturb the colchicine-dependent regulatory mechanism of Pyrin. This is supported by studies showing that Pyrin variants containing FMF mutations are resistant to inhibition by colchicine and that dephosphorylation is sufficient to activate the FMF form of Pyrin [14, 15]. However, as the B30.2 domain has been lost in the mouse, it is difficult to model this disease. Thus, there is a need for further research into the effects of these mutations in human cell types to understand how they alter Pyrin regulation.

*C. difficile* infection is a leading cause of hospital-associated mortality through diarrhea triggered by antibiotic therapy-mediated dysbiosis and pseudomembranous colitis. All of these effects are entirely dependent on the expression of TcdA and TcdB, although TcdB is sufficient to cause disease [16]. Yet, the role of the inflammasome in the immune response to *C. difficile* infection is controversial. *In vivo* experiments in ASC KO mice examining the effect of TcdA and TcdB directly, rather than through *C. difficile* infection, demonstrated that inflammasome activation in response to these toxins increased tissue damage in an IL-1β dependent manner [17]. However, a subsequent study demonstrated a role for caspase-1 in controlling *C. difficile* infection [18]. Experiments in both NLRP3 KO and Pyrin KO mice found that neither sensor impacted the severity of the disease [18, 19]. It is conceivable that the pathology of the human pathogen *C. difficile* is not fully recapitulated in the murine model. Therefore, investigations into the inflammasome response to *C. difficile* in human cells are very important, and have so far included PBMCs, monocytes, and neutrophils [11].

In this study, we assessed the inflammasome response to *C. difficile* and its toxins, TcdA and TcdB, in a human macrophage model using M-CSF monocyte-derived macrophages. We discovered that only TcdB triggered an inflammasome response in these cells and that, in contrast to monocytes, this was entirely independent of Pyrin and instead occurred through NLRP3. Furthermore, we found that prolonged exposure to LPS or type-I or –II interferons was sufficient to reactivate the Pyrin inflammasome by increasing Pyrin expression. These results demonstrate that Pyrin is held in an inactive state when monocytes differentiate to macrophages by a hitherto uncharacterized regulatory mechanism.

## Results

Inflammasome activation by *C. difficile* in hMDM is dependent on the expression of its toxins and can be blocked by NLRP3 inhibition

To investigate whether *C. difficile* or its secreted virulence factors could elicit an inflammasome response in primary hMDM, we treated the cells with either conditioned supernatant from *C. difficile* or infected hMDM with the bacteria directly. As production of TcdA and TcdB is crucial for both *C. difficile*-driven pathology and the activation of the Pyrin inflammasome, two different strains of *C. difficile* were used: one proficient for production of both toxins A and B, and one deficient for both. We also assessed the effect of prior exposure to TLR ligands on the inflammasome response by priming one hMDM group with LPS before exposure to the supernatant or bacteria. In addition, we pre-incubated one group of cells with the potent and highly specific NLRP3 inhibitor CP-456,773 (also known as CRID3 or MCC950) [20] to determine if the response was NLRP3 dependent.

We determined that both *C. difficile* conditioned supernatant and direct infection induced the release of two inflammasome-dependent cytokines, IL-1β and IL-18, from hMDM. The cytokine release was entirely toxin-dependent in the cells treated with the conditioned supernatant (Fig. 1a) and primarily reliant on toxin expression when the cells were infected directly (Fig. S1a). Notably, IL-18, which is transcribed independent of TLR stimulation [21], was still secreted exclusively when the cells were pretreated with LPS. The secretion of both cytokines was inhibited by CP-456,773 [20]. Thus, both the requirement for pre-priming and sensitivity to CP-456,773 suggested that NLRP3 mediated the response to *C. difficile* in hMDM.

**Figure 1.**
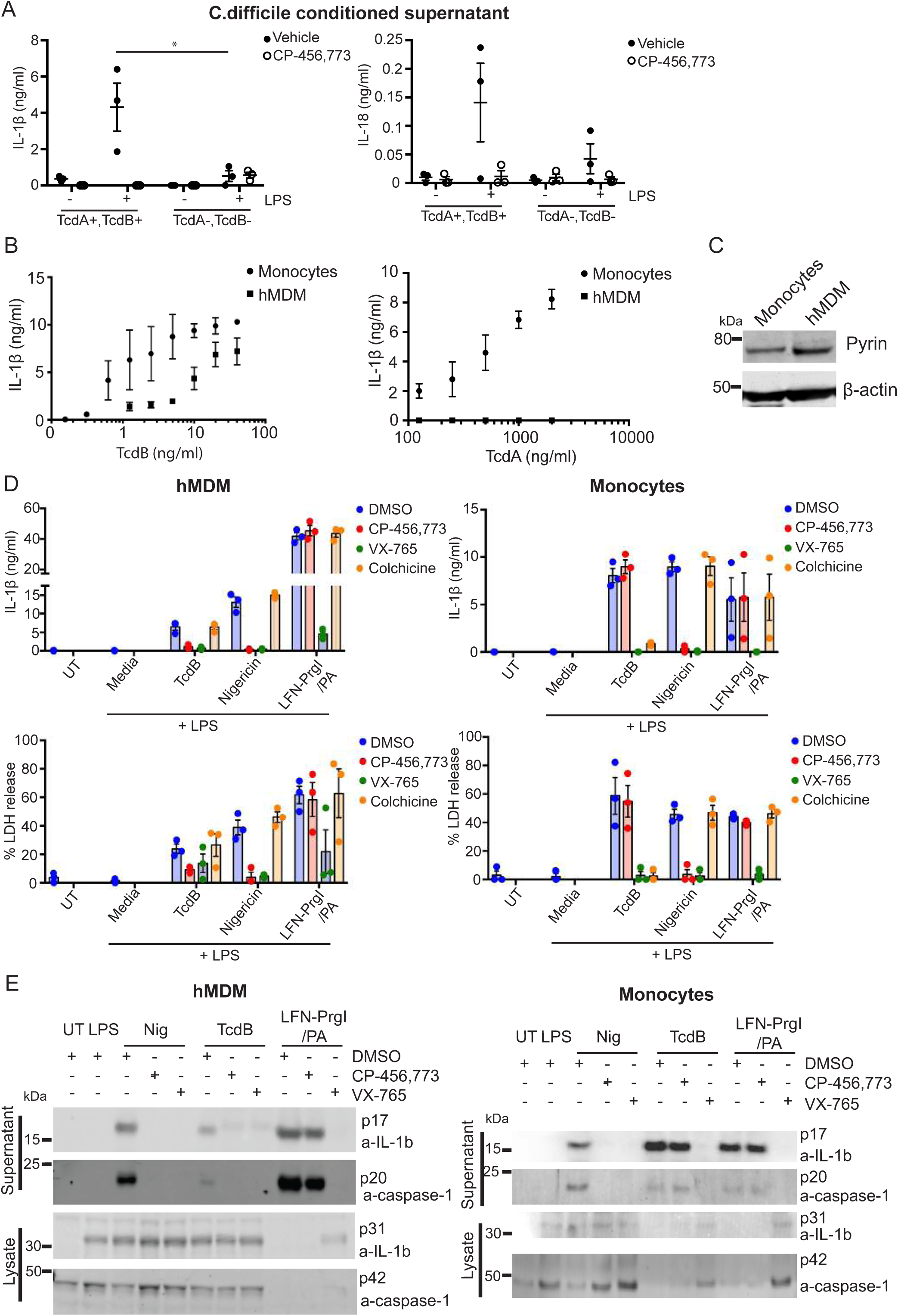
TcdB, but not TcdA, triggers CP-456,773 sensitive IL-1β release in hMDM but not monocytes. (A) hMDM incubated with conditioned supernatant from toxin proficient (TcdA+,TcdB+) or toxin deficient (TcdA-,TcdB-) *C. difficile* for 4h, then the supernatant assessed for IL-1β and IL-18. (B) LPS primed (10ng/ml, 3h) monocytes or hMDM were treated with a dose titration of either TcdB or TcdA for 3h. Supernatants were harvested and assessed for IL-1β. (C) Immunoblot of lysate from LPS treated (10ng/ml, 3h) monocytes or hMDM probed for either Pyrin or actin. Representative of 3 independent experiments. (D) LPS primed (10ng/ml, 3h) monocytes or hMDM were pre-incubated for 15 min with the vehicle alone, CP-456,773 (2.5 μM), VX-765 (40 μM) or colchicine (2.5 μM) then stimulated with TcdB (20 ng/ml), Nigericin (8 μM) or LFN-PrgI/PA (25 ng/ml) for 2.5h. The supernatant was assessed for IL-1b and LDH activity or (E) harvested and precipitated and the cells lysed. Both the precipitated supernatant and cell lysate were analysed by immunoblot for IL-1β and caspase-1. Mean and SEM shown for three donors or immunoblots representative of 3 independent experiments.

### *C. difficile* toxin B, but not toxin A, mediates NLRP3-dependent inflammasome activation in hMDM

We observered that hMDM released IL-1β and IL-18 only in response to the toxin-proficient bacteria. To determine whether one or both toxins could trigger an inflammasome response, we incubated the cells with either recombinant toxin A (TcdA) or toxin B (TcdB). As a control, we incubated monocytes with both toxins, as they have previously been shown to respond to both in a Pyrin inflammasome-dependent manner. Monocytes, as predicted, released IL-1β in response to both TcdA and TcdB (Fig. 1b). Surprisingly, and in contrast to the monocytes, we found that TcdB, but not TcdA, induced an inflammasome response in the hMDM (Fig. 1b). Notably, TcdB triggered IL-1β release from hMDM at concentrations as low as 1 ng/ml. To ensure that the lack of TcdA-mediated inflammasome activation was not due to a failure of toxin uptake, we assessed TcdA mediated Rac modification. To do so, we used a previously described monoclonal antibody that no longer recognizes Rac when its epitope is modified by the toxin [22]. The antibody was unable to bind to Rac in both monocytes and hMDM treated with the active forms of TcdB and TcdA, but not with mutants lacking glucosyltransferase activity, indicating that the cells took up the toxin (Fig. S1b).

Given the disparity in inflammasome response between monocytes and hMDM, we assessed whether Pyrin was differentially expressed in the two cell types. Pyrin levels were comparable in hMDM and monocytes (Fig. 1c), indicating that Pyrin expression was not limiting.

To further investigate the TcdB-mediated inflammasome response, we pre-incubated LPS primed hMDM or monocytes with inhibitors against NLRP3 (CP-456,773), Pyrin (colchicine) or caspase-1 (VX-765) and then treated the cells with TcdB. As specificity controls, we used Nigericin, a potassium ionophore that activates NLRP3, as well as the NLRC4 activator needletox. The latter contains the *Salmonella typhimurium* T3SS needle protein PrgI fused to the N-terminus of anthrax lethal factor, which is delivered to the cytosol by anthrax protective antigen (PA). In hMDM, we determined that both TcdB dependent IL-1β secretion and LDH release as a measure of pyroptosis were inhibited by CP-456,773 but not colchicine. This provides further evidence that TcdB is activating NLRP3 rather than Pyrin in these cells (Fig. 1d). In contrast, and as demonstrated previously, the response to TcdB in monocytes was not inhibited by CP-456,773 but was inhibited by colchicine, indicating a dependence on Pyrin (Fig. 1d). To ensure that the observed IL-1β release was accompanied by caspase-1 activation, we also assessed the cleavage of IL-1β and caspase-1. In agreement with the IL-1b and LDH release results, we found that TcdB-mediated IL-1β and caspase-1 cleavage were CP-456,773 sensitive in hMDM, but not in monocytes (Fig. 1e). We also monitored the actin cytoskeleton after treatment by TcdB by staining the cells with Phalloidin. As predicted, TcdB disrupted the actin cytoskeleton similarly in both cell types (Fig. S1c).

### The inflammasome response to TcdB in human macrophage cell lines is entirely dependent on NLRP3

Our results thus far demonstrated that NLRP3 is the responding inflammasome sensor to TcdB in hMDM, but relied solely on compound-based inhibition. Therefore, we used genetically modified human macrophage cell lines to determine whether NLRP3 or Pyrin mediates the inflammasome response to TcdB. Initially, we sought a model cell line that had a robust inflammasome response to TcdB. We investigated whether the BLaER1 cell line, a recently established monocyte/macrophage cell line [23], would similarly respond to TcdB to the hMDM. We found that WT BLaER1 cells released IL-1β in response to treatment with TcdB, as well as activating caspase-1 as measured by a caspase-1 activity assay (Fig. 2a). These data demonstrate that TcdB activates an inflammasome response in BLaER1 similar to hMDM.

**Figure 2.**
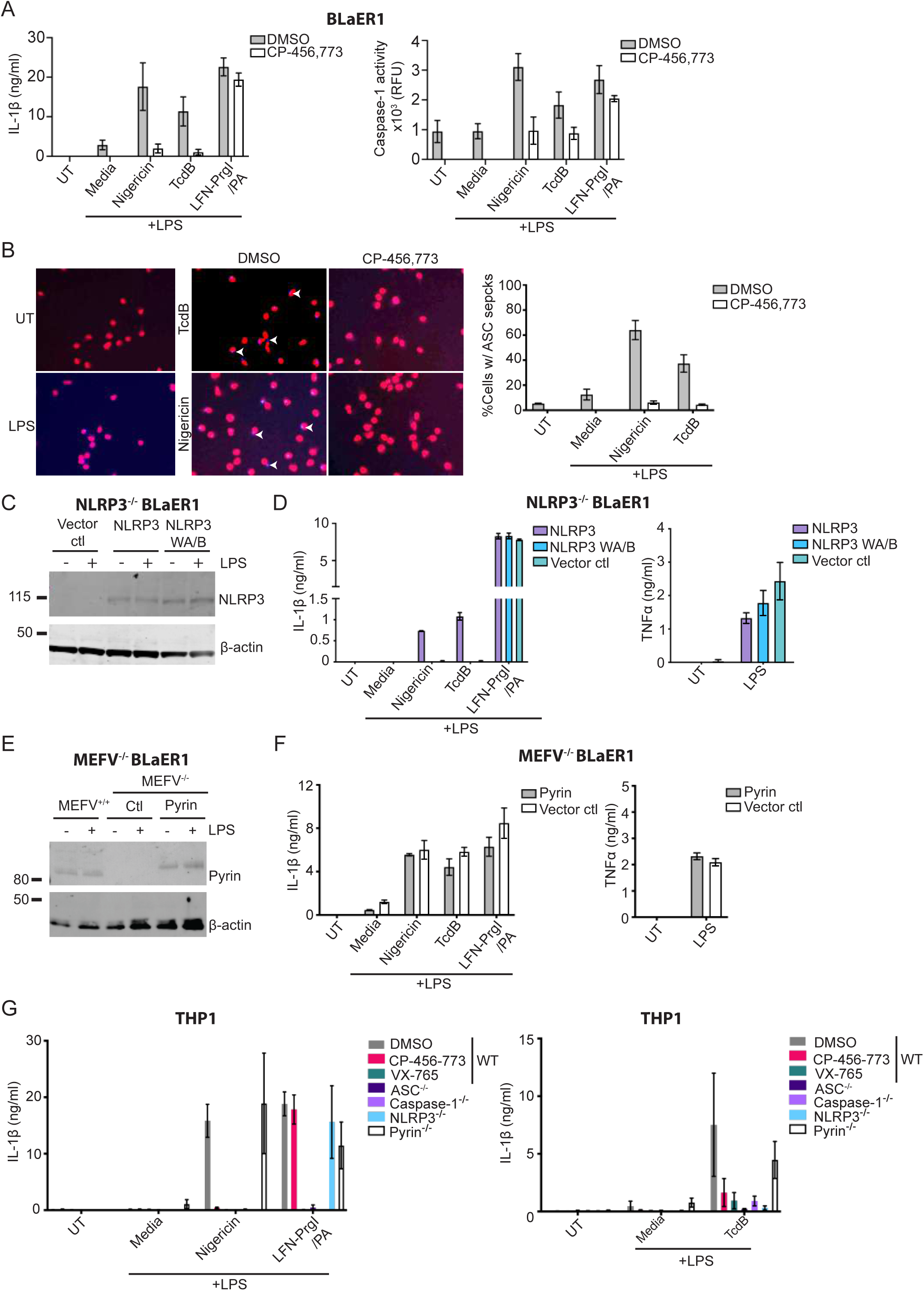
NLRP3, not Pyrin, is the responding inflammasome sensor to TcdB in human macrophages. (A) Differentiated WT BLaER1 cells were primed with LPS (100 ng/ml, 3h), pre-incubated with either CP-456,773 (2.5 μM, 15 min), then activated with Nigericin (8 μM), TcdB (20ng/ml) or with LFN-PrgI/PA (25 ng/ml each) for 2h. IL-1β and caspase-1 activity were assessed from the harvested supernatants. (B) ASC-BFP transduced WT BLaER1 cells treated as in (A). Cells were then fixed and the number of ASC specks quantified by microscopy. (C) NLRP3 expression in differentiated BLaER1 cells (+/-100ng/ml LPS 3h) was assessed by immunoblot. (D) Differentiated Casapse-4, NLRP3 double deficient BLaER1 cells reconstituted with either NLRP3, the NLRP3 walker A/B mutant (NLRP3 WAB) or the vector control treated as in (A) and the supernatants assessed for IL-1β or TNFα. Mean and SD of three technical replicates shown, representative of 3 independent experiments. (E) Immunoblot of Pyrin expression in differentiated BLaER1 cells. Representative of 3 independent experiments. (F) Differentiated Pyrin deficient BLaER1 cells reconstituted with either Pyrin or the vector control treated as in (A) and the supernatants assessed for IL-1β or TNFα. Mean and SD of three technical replicates shown, representative of 3 independent experiments. (G) LPS primed WT THP-1s or the listed KOs were activated with inflammasome activators for either 1.5h (Nigericin, LFN-PrgI/PA) or for 8h (TcdB). Supernatants were assessed for IL-1β. Where used, CP-456,773 (2.5 μM) and VX-765 (40 μM) were pre-incubated with the cells for 15 min prior to addition of the inflammasome activators. Mean and SEM of 3 independent experiments shown.

We next used the BLaER1 cell line to determine the propensity of TcdB to trigger ASC speck formation, another hallmark of inflammasome activation. Accordingly, we generated a BLaER1 cell line overexpressing ASC-mCherry and stimulated it with either TcdB or Nigericin in the presence or absence of CP-456,773. This experiment was also performed in the presence of the caspase-1 inhibitor VX-765 to prevent cell death of inflammasome-activated cells. Following stimulation, the cells were fixed, and the number of ASC specks was quantified by microscopy, followed by normalization to the number of nuclei in each image. Consistent with our other data, we found that TcdB and Nigericin also caused CP456,773-sensitive ASC speck formation in these cells (Fig. 2b).

To determine the sensor responsible for the TcdB mediated inflammasome response, we used either NLRP3 or Pyrin knock-out BLaER1 cells. As the NLRP3 KO cells we used were on a caspase-4 KO background, we first ensured that caspase-4 played no role in the response. Accordingly, we tested the inflammasome response to TcdB in the caspase-4 KO BLaER1 cells, but found no difference in IL-1β secretion, demonstrating caspase-4 was not required for the response to TcdB (Fig. S2a). The NLRP3, caspase-4 double KO cells were then reconstituted with either NLRP3, the inactive Walker A/B NLRP3 mutant, or a vector alone control. NLRP3 expression was confirmed by immunoblot (Fig. 2c). As previously, we primed these cells with LPS, incubated them with either TcdB, Nigericin or needletox, and assessed IL-1β release to the supernatant. In agreement with our findings in hMDM, only the cells reconstituted with active NLRP3, but not the walker A/B mutant, were able to respond to TcdB and Nigericin (Fig. 2d). In contrast, all cell lines responded equally to the NLRC4 trigger needletox (Fig. 2d) and secreted similar levels of TNFa in response to LPS (Fig. 2d). These results confirm that the TcdB-mediated inflammasome response was dependent on the expression of active NLRP3.

To confirm that Pyrin was not required for the TcdB-driven inflammasome response, we reconstituted Pyrin KO BLaER1 cells with either Pyrin or vector control, and confirmed expression levels by immunoblot (Fig. 2e). We found the inflammasome response to TcdB was unaffected by the absence of Pyrin, indicating that Pyrin does not play a role in the inflammasome response to TcdB in these cells (Fig. 2f). Similarly, Pyrin expression did not affect the inflammasome response to Nigericin or needletox (Fig. 2f).

BLaER1 cells can readily secrete IL-1β in response to TLR4 stimulation by LPS alone. To rule out that LPS contamination of the TcdB was responsible for the observed IL-1β release, we stimulated the cells with TcdB or Nigericin following pre-incubation of the cells with TAK-242, a TLR4 inhibitor. TAK-242 did not block TcdB-mediated IL-1β release (Fig. S2b), but TAK-242 completely abolished LPS mediated TNFa secretion when applied before LPS stimulation (Fig. S2c), demonstrating that NLRP3 mediated activation by TcdB is not due to LPS contamination or TLR4 activation.

Having established the requirement for NLRP3 in response to TcdB in the BLaER1 cell line, we sought to determine if this was true in the other commonly used human macrophage cell line, THP-1. We first titrated TcdB on PMA differentiated THP-1 cells and found that it required both a higher concentration of TcdB (2ug/ml) to trigger IL-1β release as well as a longer incubation time (8h). Having established this we tested several THP-1 CRISPR knock-out cell lines to determine the requirements for TcdB-mediated inflammasome activation. PMA-differentiated WT THP-1 cells or THP-1 cells deficient in ASC, caspase-1, NLRP3 or Pyrin were primed with LPS (200 ng/ml, 3h), then incubated with TcdB, Nigericin and needletox. WT cells were also pre-incubated with CP-456,773 to determine if it had the same effect as ablation of NLRP3. Echoing our previous data, the TcdB-triggered inflammasome response in THP-1 cells was CP-456,773-sensitive and NLRP3-dependent (Fig. 2g). As anticipated, this response also required ASC and caspase-1, but not Pyrin. Nigericin similarly was NLRP3-dependent, while the NLRC4 trigger only required ASC and caspase-1, demonstrating the effect of NLRP3 ablation was specific (Fig. 2g). Furthermore, the different KO lines secreted similar amounts of TNFα in response to LPS (Supp. Fig 2d). Collectively, these results show an absolute requirement for NLRP3, but not Pyrin, in the TcdB-mediated inflammasome response in human THP-1 cells.

Having established that TcdB activates NLRP3, but not Pyrin, we next determined whether NLRP3 activation required the activity of the TcdB glucosyltransferase domain (GTD) against Rho, as found for Pyrin. Thus, we incubated LPS-primed hMDM with TcdB or a variant of TcdB containing inactivating mutations in the glucosyltransferase domain, D286N and D288N (TcdB NXN), which does not inactivate Rho, Rac or Cdc42. We determined that both TcdB and TcdB NXN induced IL-1β release (Supp. fig. 2e). Interestingly, the NXN variant of TcdB was a more proficient inflammasome activator than the WT toxin, suggesting that the activity of the GTD domain opposes TcdB-mediated NLRP3 activation. This demonstrates that in contrast to Pyrin activation, GTD activity, and subsequent Rho inhibition are not required for the TcdB mediated inflammasome response in hMDM.

### TcdA and TcdB elicit an NLRP3-independent inflammasome response in BMDM and peritoneal macrophages

The observation that neither TcdB nor TcdA could activate Pyrin in hMDM was unexpected. To test if this was specific to human cells, we next analyzed toxin-dependent inflammasome responses in differentiated macrophages from bone marrow (BMDM) as well as isolated peritoneal macrophages (PM) from either WT or NLRP3 KO mice. We stimulated them with either TcdA or TcdB, using Nigericin and poly(dA:dT) transfection as specificity controls for NLRP3 and AIM2, respectively.

We determined that TcdA and TcdB triggered IL-1β secretion in BMDM from both WT and NLRP3 KO cells, indicating that the response was NLRP3 independent (Fig. 3a). This was also true in peritoneal macrophages, which showed little difference in toxin-mediated IL-1β secretion between WT and NLRP3 KO cells (Fig. 3b). In contrast, the inflammasome response to Nigericin was ablated entirely in the NLRP3 KO. At the same time, there was no difference in IL-1β release in response to transfected dA:dT and no difference in TNFa secretion in response to LPS. We also assessed IL-1β and caspase-1 cleavage in WT and NLRP3 KO BMDM following LPS priming and stimulation with TcdB, Nigericin or dA:dT. Similarly, we found no differences between the two genotypes when stimulated with TcdB or the specificity control dA:dT, while Nigericin mediated IL-1β and caspase-1 cleavage were ablated in the NLRP3 KO (Fig. 3c).

**Figure 3.**
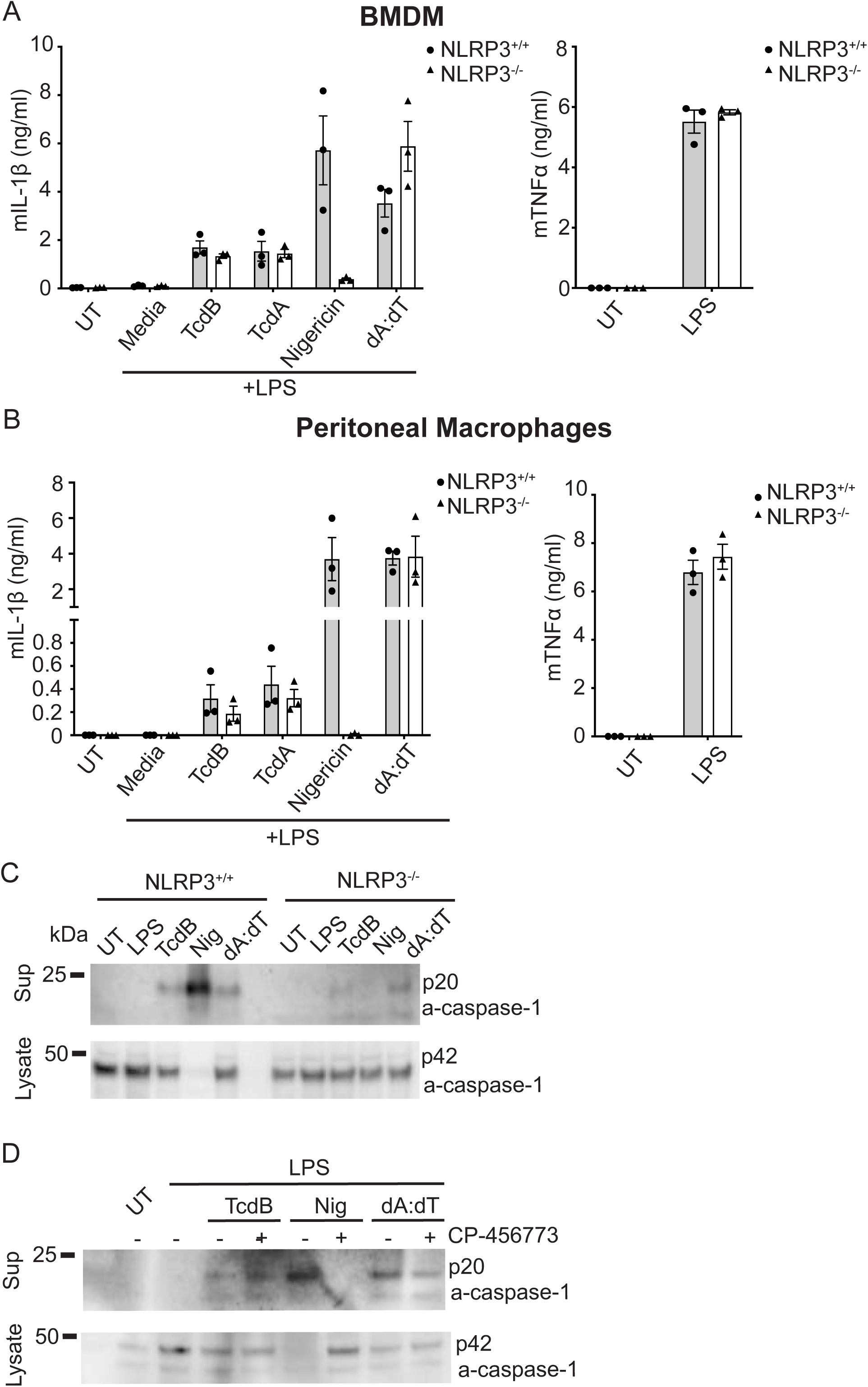
Pyrin is the responding sensor to TcdB in murine macrophages. WT and NLRP3 deficient BMDM (A) or peritoneal macrophages (PM) (B) were primed with LPS (200 ng/ml, 3h), then activated with Nigericin and TcdB for 2h or dA:dT for 4h. (C) IL-1β and caspase-1 immunoblot of precipitated supernatant or cell lysate from WT and NLRP3 deficient BMDM treated as in (A). (D) IL-1β and caspase-1 immunoblot of precipitated supernatant or cell lysate from LPS primed WT BMDM either untreated or pretreated with CP-456,773 (2.5 μM, 30 mins), then stimulated as in (A). Mean and SEM of 3 independent experiments shown, immunoblots are representative of 3 independent experiments.

Given that we rely on CP-456,773 to determine the role of NLRP3 in response to TcdB in human primary macrophages, we also investigated whether, despite the lack of NLRP3 dependence, CP-456,773 could affect the response in BMDM. Therefore, WT BMDM were primed with LPS pre-incubated in the presence or absence of CP-456,773 and then incubated with TcdB, Nigericin and dA:dT. However, as with the NLRP3 KO cells, CP-456,773 did not affect TcdB- or dA:dT-mediated IL-1β or caspase-1 cleavage but completely inhibited Nigericin-mediated cleavage of both (Fig. 3d). Therefore, the TcdB-mediated inflammasome response in murine macrophages is both NLRP3-independent and insensitive to CP-456,773. Collectively these results show that the inflammasome response to both TcdA and TcdB in murine macrophages is independent of NLRP3 and likely depends on Pyrin.

### Prolonged incubation with LPS or type I and II interferons increases Pyrin expression in hMDM

It was surprising that, in contrast to monocytes, neither TcdA nor TcdB triggered a Pyrin inflammasome response in hMDM. Thus, we next investigated whether inflammatory conditions increased the expression of inflammatory signaling molecules and thus potentially enable their activation. Pro-inflammatory signaling molecules activating either the NF-κB or IRF transcription factors have been demonstrated to increase Pyrin expression in PBMCs [24]. To determine whether either of these pathways could increase Pyrin expression in hMDM, we treated the cells with LPS, Pam3CS4K, TNFα, IFN-β and IFN-γ, as well as IL-4 and IL-10 for 5 or 18 hours and assessed Pyrin expression by immunoblot. Notably, only LPS increased Pyrin expression after 5h, while LPS, IFN-β and IFN-γ but not TNFα, IL-1β or Pam3CS4K increased expression after 18h (Fig. 4a). Thus, only stimuli that signal through IRF family transcription factors triggered an increase in Pyrin expression. To ensure that these molecules were functional in our system, we assessed the expression of IL-1β, a target of NF-kB, in response to LPS and Pam3CS4K. We determined that both were able to increase IL-1β expression (Fig. 4a), demonstrating that they were functional but unable to cause an increase in Pyrin expression.

**Figure 4.**
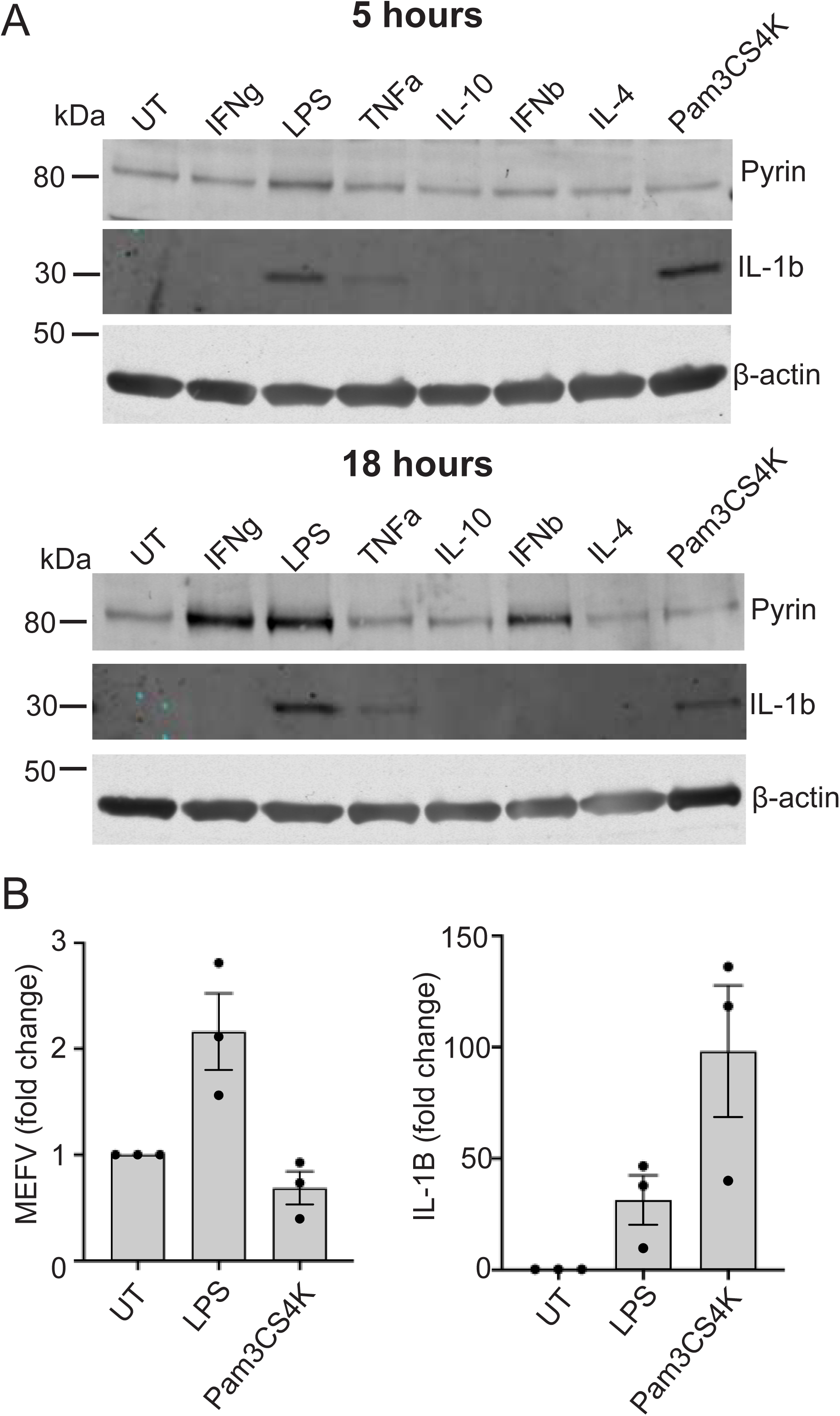
LPS type I and type II interferon increase Pyrin expression and enable Pyrin activation in human macrophages. (A) Pyrin and IL-1β expression in hMDM treated with either IFN-γ (200 U/ml), LPS (10ng/ml), TNFα(50ng/ml), IL-10 (100ng/ml), IFN-β (5000 U/ml), IL-4 (1000U/ml) or Pam3Cys4K (20ng/ml) for either 5 or 18h. Representative of 3 independent experiments. (B) Pyrin (MEFV) or IL-1β transcript from hMDM treated LPS (10ng/ml) or Pam3Cys4K (20ng/ml) for 12h. Mean and SEM the fold change of three experimental replicates shown.

Previous studies had shown that the Pyrin promoter contains an IRSE element that can be activated by both TRIF and IFN signaling [24], suggesting the increase in Pyrin expression observed can be due to increased transcription. We investigated this by stimulating the cells with either LPS or Pam3 for 12h, then assessing mRNA copy number for Pyrin by qPCR, using IL-1β as a control. We observed that LPS, but not Pam3CS4K, caused an increase in *MEFV* transcript compared to the untreated cells (Fig. 4b). In contrast, both LPS and Pam3CS4K increased IL-1β transcription, demonstrating that the increase in *MEFV* transcript was specific to LPS (Fig. 4b). It is, therefore, likely that the increase in Pyrin expression is driven by increased gene transcription.

### LPS and interferons prime activation of the Pyrin inflammasome in hMDM

We tested whether increased Pyrin expression would be sufficient to enable Pyrin inflammasome activation. Accordingly, we primed hMDM with LPS for either 3 or 18 hours, pre-incubated them with DMSO, CP-456,773, VX-765 or colchicine, and then treated them with TcdA or Nigericin. As we had observed previously, TcdA did not trigger an inflammasome response after 3h of LPS priming (Fig. 5a). In contrast, after 18 h, TcdA triggered robust secretion of IL-1β that was sensitive to colchicine, and thus dependent on Pyrin (Fig. 5a). By comparison, Nigericin mediated CP-456,773-sensitive IL-1β release after both 3 and 18 hours of LPS priming but was not affected by colchicine (Fig. 5a). Similarly, priming the cells for 18h, but not 3h, with IFN-β was sufficient to trigger Pyrin-dependent IL-18 secretion, demonstrating that it could also prime a Pyrin response (Fig. 5b).

**Figure 5.**
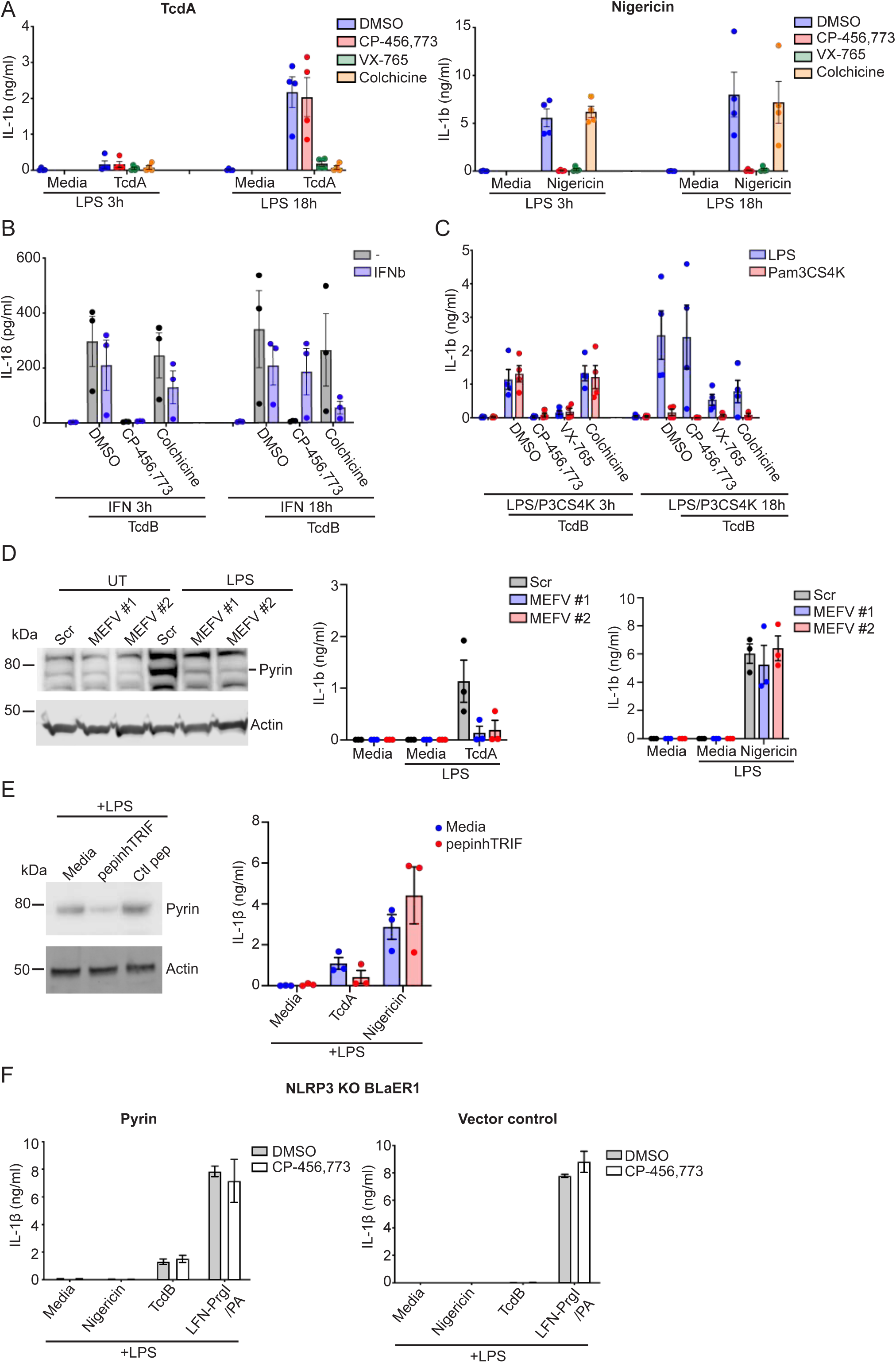
Increased Pyrin expression is necessary and sufficient for Pyrin reactivation in human macrophages. (A) LPS primed (10ng/ml, 3 or 18h) hMDM were pre-incubated with compounds as noted previously, then stimulated with either TcdA (200 ng/ml) or Nigericin (8 μM) for 2.5h. The supernatant was harvested and assessed for IL-1β release. (B) IFN-β(5000U/ml, 3 or 18h) primed hMDM with either for 3h or 18h were pre-incubated with compounds as before then stimulated with TcdB (20 ng/ml) for 2.5h. The supernatant was harvested and assessed for IL-18 release. (C) hMDM primed with either LPS (10ng/ml) or Pam3Cys4K (20ng/ml) for 3h or 18h were treated as in (B). The supernatant was harvested and assessed for IL-1β release. (D) hMDM were transfected with siRNA targeting Pyrin mRNA (#59, #60) or the scrambled control and assessed for Pyrin expression by immunoblot or stimulated a previously following primed with LPS (10ng/ml) for 18h. (E) Expression of Pyrin in hMDM incubated with pepinhTRIF or the scrambled control peptide (25μg/ml, 5h), then treated with LPS (10ng/ml, 18h). Representative of 3 donors. IL-1β secretion from cell pretreated with pepinhTRIF or the scrambled control peptide (25μg/ml) in response to LPS priming (10ng/ml, 18h) followed by stimulation with either TcdA (200ng/ml) or nigericin (8μM) for 2.5h. (F) LPS primed (100ng/ml, 3h) differentiated Casapse-4, NLRP3 double deficient BLaER1 cells reconstituted with either Pyrin or the vector control were incubated with Nigericin (8μM) TcdB (20ng/ml) or LFN-PrgI in the presence of protective antigen (PA) (25ng/ml) for 2.5h in the presence or absence of CP-456,773 (2.5μM, 15 min incubation). Supernatants were assessed for IL-1β. For (A)-(E) Mean and SEM of 3-4 experimental replicates shown. For (F) Mean and SD of three technical replicates shown, representative of 3 independent experiments.

We next sought to determine whether restoration of the Pyrin inflammasome response was specific to a stimulus that increased Pyrin expression. We thus compared the ability of TcdB to activate Pyrin in hMDM primed with Pam3CS4K to those primed with LPS. As done previously, hMDM were pre-incubated with different inhibitors to determine the responding inflammasome sensor. TcdB triggered CP-456,773-dependent inflammasome activation after three hours of priming with either LPS or Pam3CS4K (Fig. 5c). Comparative stimulation after 18 hours of priming led to a Pyrin-dependent response in the LPS primed cells, while the Pam3CS4K-primed cells failed to activate any inflammasome response (Fig. 5c), demonstrating that increased Pyrin expression is linked to Pyrin reactivation.

### Increased Pyrin expression is necessary for the Pyrin response in hMDM

To determine whether the increase in Pyrin expression was a requirement for Pyrin activation, we transfected hMDM with two distinct siRNA against Pyrin or a scrambled control 24h before priming with LPS. Notably, both Pyrin targeting siRNAs effectively prevented the LPS mediated increase in Pyrin expression. Still, they did not reduce it further than the untreated control, while the scrambled control had no effect (Fig. 5d). siRNA transfected hMDM were primed with LPS for 18h and then stimulated with TcdA or Nigericin. We observed that the two Pyrin siRNAs, but not the scrambled control, prevented TcdA-mediated IL-1β release. In contrast, neither of the Pyrin-targeting siRNAs, nor the control siRNA, affected Nigericin-mediated inflammasome activation (Fig. 5d), establishing decreasing Pyrin expression is sufficient to inhibit Pyrin activation specifically.

As Pyrin expression increased after treatement with LPS or interferons, but not Pam3Cys4K or TNFα, we hypothesised that the increase in LPS-dependent Pyrin expression likely required the TRIF signaling pathway. We tested our hypothesis by investigating whether blocking TLR4-mediated TRIF signaling prevents the LPS-dependent increase in Pyrin expression and restores the Pyrin inflammasome response. To this end, we pretreated hMDM with pepinhTRIF, a peptide that prevents the interaction of TRIF with its downstream interaction partners. We then incubated these cells with LPS for 18h before assessing Pyrin expression and activation. We found that treatment with pepinhTRIF, but not a control peptide, reduced LPS mediated Pyrin expression (Fig. 5e). We then stimulated the cells with either TcdA or Nigericin. We found that only the TcdA-mediated inflammasome response was inhibited by pretreatment with the pepinhTRIF, whereas the Nigericin-mediated IL-1β secretion was unaffected (Fig. 5e). These results demonstrate that the LPS-stimulated increase in Pyrin expression is TRIF-mediated, and blocking this is sufficient to prevent Pyrin activation in these cells.

Given that increased Pyrin expression is necessary for Pyrin reactivation in hMDM, we next sought to determine whether Pyrin overexpression alone would enable a Pyrin inflammasome response. For this experiment we used the caspase-4, NLRP3 KO BLaER1 cells, which otherwise do not mount an inflammasome response to TcdB. We overexpressed Pyrin in the caspase-4, NLRP3 KO BLaER1 cells using reconstitution with a vector alone as a control. We then primed these cells with LPS and stimulated them with TcdB, Nigericin or needletox. Markedly, TcdB mediated an inflammasome response in the Pyrin-reconstituted cells, but not in cells transduced with the vector alone Notably, this was not inhibited by CP-456,773 (Fig. 5f). In contrast, Pyrin reconstitution had no effect on either NLRP3 or NLRC4 activation (Fig. 5f). This differed from our earlier observations where TcdB triggered an NLRP3 dependent inflammasome response in Pyrin KO BLaER1 cells overexpressing Pyrin. However, in this experiment we did not determine whether Pyrin overexpression in the Pyrin KO BLaER1 cells resulted in NLRP3 independent IL-1β secretion in response to TcdB. Furthermore, when NLRP3 is present, the NLRP3 inflammasome response to TcdB might predominate in BLaER1. These results demonstrate that an increasing Pyrin expression is sufficient to enable Pyrin inflammasome response, and regulation of Pyrin expression is the primary factor controlling Pyrin inflammasome activation in hMDM.

### Dephosphorylation of Pyrin is insufficient for activation in hMDM

Pyrin does not form an inflammasome in response to TcdA or TcdB in hMDM in the absence of a priming stimulus. This suggests that the toxin cannot disengage one or both of the two known inhibitory mechanisms controlling Pyrin: either dephosphorylation of the S204 and S242 residues, or the less well characterized mechanism related to the B30.2 domain. To determine the involvement of phosphorylation in regulating Pyrin activation in hMDM we tested whether TcdB mediated dephosphorylated of Pyrin at serine 242 after priming with LPS for 3h or 18h. Notably, we found that Pyrin was dephosphorylated after treatment with TcdB regardless of the length of LPS priming, demonstrating that this is unlikely to be the mechanism preventing Pyrin activation (Fig. 6).

**Figure 6.**
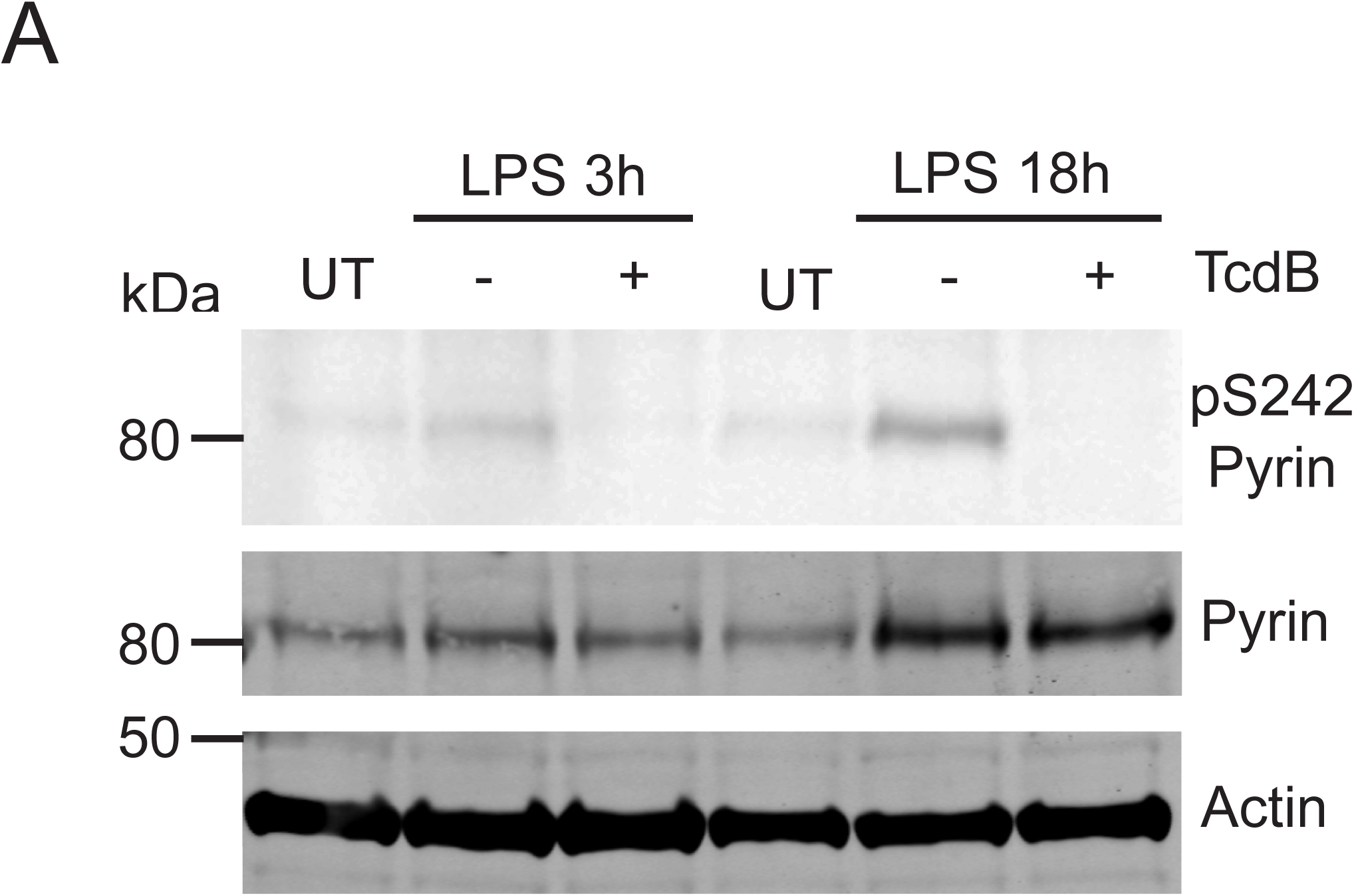
The B30.2 domain regulates Pyrin activation in human macrophages, and is disrupted by FMF mutations. (A) LPS primed (10ng/ml, 3h or 18h) hMDM were treated with TcdB (5 ng/ml, 1h), then lysed and assessed for phosphorylation of Pyrin (S242), Pyrin or actin by immunoblot. Representative of 3 independent experiments.

## Discussion

Pyrin responds to virulence factors and endogenous molecules that inhibit the RhoA signaling pathway. Gain of function mutants in the MEFV gene cause multiple genetic autoinflammatory disorders [11]. Understanding Pyrin regulation, particularly in the context of different human cell types, is therefore critical to understand how auto-inflammation-associated mutations may disrupt these mechanisms. In this study, we investigated the inflammasome response to the disease-causing *C. difficile* toxins TcdA and TcdB in hMDM. We found that under steady state conditions, and in spite of expressing similar levels of Pyrin to monocytes, hMDM do not mount a Pyrin inflammasome response. Notably, Pyrin activation could be re-established in these cells by further increasing in Pyrin expression through prolonged priming with either LPS or type I or II interferon. Surprisingly, in the absence of Pyrin activation, TcdB instead activated NLRP3 in these cells, thus demonstrating a redundancy in the inflammasome system to detect this toxin. TcdB similarly triggered NLRP3 activation in both human macrophage cell lines we tested, BLaER1 and THP1, rather than Pyrin. Notably, unlike Pyrin activation, NLRP3 activation by TcdB did not require the enzymatic activity of the toxin, demonstrating that it activates Pyrin and NLRP3 through two distinct pathways. These findings are in contrast to monocytes, which responded to both toxins in a Pyrin inflammasome dependent manner, as has been shown previously [3]. This demonstrates a cell type specific divergence in the inflammasome response to TcdB.

The results of our study demonstrate that regulatory mechanisms governing Pyrin activation in hMDM prevent Pyrin activation unless the cell has been exposed to a prior pro-inflammatory stimulus. Notably, in the case of Pyrin, licensing required prolonged stimulation with LPS or interferons, but not other inflammatory stimuli including TNFα. This contrasts with the data from a previous study in PBMCs, where TNFα and Pam3CS4K stimulation also increased Pyrin expression [24]. Increased expression of Pyrin was necessary to prime a Pyrin dependent inflammasome response in hMDM, as preventing increased Pyrin expression after these stimulation with LPS was sufficient to inhibited Pyrin activation. Furthermore, overexpression of Pyrin in BLaER1 cells was sufficient to enable a Pyrin response. These results suggest that, similar to the NLRP3 and AIM2 inflammasomes, control of Pyrin expression through inflammatory signals regulates Pyrin activation. Although the promoter region of the MEFV gene contains elements that are recognized by either interferon driven transcription factors or by NF-kB [24], increased Pyrin expression was specific to activation of the TRIF/interferon pathway. This suggests that Pyrin is specifically reactivated in hMDM by pathways associated with cell host defense and gram-negative bacteria and may be important in the response to these pathogens. An example of this is an increase in Pyrin expression seen in response to *Francisella novidica* infection, where, unlike in mice, Pyrin is the responding sensor to *F. novicida* in hMDM [25]. This also has implications for Pyrin associated auto-inflammatory conditions, where, compared to NLRP3 driven auto-inflammatory conditions, IL-1β blocking therapy is only somewhat successful in preventing disease [26]. Our data suggests that interferon signaling could also play a role in triggering auto-inflammation in these patients, as well as DAMPs released after tissue injury that stimulate TLR4. Further research is needed to understand the contribution of these sensors to Pyrin based auto-inflammatory conditions to determine if blocking these priming signals helps to control these diseases.

Notably, we found Pyrin expression was comparible between hMDM and monocytes under unstimulated conditions, indicating that the additional regulatory steps in hMDM are post-translational. The most well characterized of these is the phosphorylation of Pyrin. However, in spite of failing to trigger formation of a Pyrin inflammasome, TcdB still caused dephosphorylation of the S242 residue, demonstrating that this is not the mechanism preventing Pyrin activation. An alternative mechanism controlling Pyrin activation could be the control mechanism related to the colchicine sensitive requirement for Pyrin activation, which requires the B30.2 domain [15]. Supporting this is the finding that, unlike in human macrophages, TcdA and TcdB activate Pyrin in murine macrophages, where a frame shift mutation has removed the B30.2 domain. Another option would include the protein proline serine threonine phosphatase-interacting protein 1 (PSTPIP1), which binds to Pyrin and facilitates its oligomerisation and inflammasome formation [27]. However, further investigations into how this control mechanism functions are required to gain further information into whether this is indeed the control mechanism in macrophages.

These findings demonstrate yet another point of divergence between human monocytes and macrophages, as Pyrin is inactive in these cells. These observations join a growing list of differences in inflammasome activation between these cell types, which also differ in their responses to LPS, which activates NLRP3 in monocytes without a need for a second stimulus, as well as dsDNA, which triggers a STING- and NLRP3-dependent inflammasome response in monocytes rather than activating AIM2 [23, 28]. That Pyrin is active in monocytes is unsurprising, as they are migratory cells that rely on actin rearrangement to reach sites of infection where they contribute to the immune response and clearing the pathogen. Therefore, inhibition of migration represents a disruption of a basic function of these cells. This is consistent with Pyrin activation in neutrophils, another migratory immune cell type. Conversely, our results demonstrate that Pyrin is unable to respond in hMDM, even though inactivation of Rho will also impact the immune response and viral clearance. This suggests a requirement for more nuanced control of Pyrin activation in hMDM, which may be a requirement to limit Pyrin driven auto-inflammation. Limiting the Pyrin response to situations requiring pre-exposure to pro-inflammatory molecules such as LPS or interferons may be necessary to ensure that Pyrin is not aberrantly activated by endogenous molecules such as bile acids [29], which may be present in tissues but not in the circulation [30]. How widespread this mechanism is will be elucidated by research focusing on Pyrin regulation in other cell types. Notably, the regulation of Pyrin observed in human macrophages was not evident in murine macrophages, which, similar to human monocytes, responded to TcdA and TcdB in a Pyrin-dependent manner. However, murine Pyrin lacks the B30.2 domain, and so may have lost the regulatory mechanism preventing Pyrin activation in hMDM. This demonstrates a further divergence in inflammasome responses between the two species, in addition to NLRC4 activation and NLRP3 responses [31, 32].

It was interesting to note that Pyrin activation was restricted in the hMDM in spite of similar basal Pyrin expression to monocytes. The expression observed was most likely due to M-CSF, which increases Pyrin expression in monocytes/macrophages [33]. However, it is unclear why Pyrin is present but stringently regulated. One possible explanation for this is that Pyrin has additional roles in the cell aside from forming an inflammasome. It has been suggested previously that Pyrin operates as a specialized adapter for autophagic machinery [34]. This capacity it associates with autophagic adapters ULK1 and Beclin1 to target NLRP3 and capase-1 for autophagic degradation. Further investigation would be required to understand how M-CSF driven Pyrin expression controls this phenomenon.

Our results have potential implications for FMF pathogenesis. Whilst defining which mutations can cause FMF is still an area of ongoing research, the most well characterized FMF-causing mutations occur in the B30.2 domain [35]. If indeed the mechanism holding Pyrin inactive requires the less well characterized mechanism involving the B30.2 domain, our data suggest that this control mechanism will be disrupted by these mutations. Further research investigating the effect of FMF mutations on regulation of Pyrin in macrophage will elucidate whether these mutations indeed alter this regulation.

Our findings have implications for the pathogenesis of *C. difficile* infection. TcdB triggered either a Pyrin- or NLRP3-dependent inflammasome response depending on the cell type, while TcdA did not trigger an inflammasome response in macrophages. This redundancy is quite intriguing, and suggests that inflammasome-mediated detection of TcdB is important in the response to the bacteria. Given that strains of *C.difficile* exist that express only one of TcdA or TcdB, as well as both [36], it is also possible that disease severity alters depending on whether the bacteria express TcdB, as early NLRP3 activation in the macrophages prior to systemic penetration by the toxin may dictate the speed of the immune response or conversely enhance tissue damage. Furthermore, the detection mechanism for each sensor has different requirements, as Pyrin detects the activity of the glycosyltransferase domain through RhoA inactivation, whilst NLRP3 mediated TcdB detection is independent of glycosyltransferase activity. Furthermore, our data demonstrate that Pyrin activation is differentially regulated in human and mouse, and so the inflammasome response to this infection may differ from what has been shown in mouse models. Further studies in a model more closely related to humans, such as pigs, is needed to determine the role of inflammasomes in *C.difficile* infection, and determine whether the current inflammasome inhibitors being developed represent new treatments to prevent *C. difficile*-associated pathology or pose an increased risk of *C. difficile* infection.

## Materials and methods

### Ethics statement

Ethics for the use of human material was obtained according to protocols accepted by the institutional review board at the University Clinic Bonn; local ethics votes Lfd. Nr. 075/14). No consent was taken as all donors were anonymous.

### Reagents

LPS (Eb-ultrapure, 0111:B4), Pam3CS4K, pepinhTRIF and TAK-242 were obtained from InvivoGen, Nigericin was obtained from Invitrogen and *Bacillus anthracis* protective antigen was obtained from List Biological Laboratories. Colchicine was obtained from Sigma, and VX-765 was obtained from Sellekchem. DRAQ5 was purchased from eBioscience. TNFα, IFN-β, IFN-γ, M-CSF, IL-3, IL-4 and IL-10 were purchased from Immunotools. TcdA and TcdB from C. difficile strain VPI10463 were recombinantly produced and supplied by Prof. Ralf Gerhard [37]. Both toxins are identical to TcdA and TcdB from strain cdi630, which was used for infection assay. The HTRF kits for human IL-1β and TNFa were obtained from Cisbio, the ELISA kit for human and mouse IL-1β was obtained from R&D Systems. Both were used according to the manufacturer’s instructions. For the *Clostridioides difficile* supernatant transfer and co-culture, the following reagents were used: Butyric acid (Sigma-Aldrich: W222119-1KG-K), Various amino acids (Roth or Sigma), Iron sulfate heptahydrate (Sigma-Aldrich: 215422-250G), Triton-X 100 (Roth: 3051.2), M-Per (Sigma-Aldrich: 78501), Protease inhibitor (Sigma-Aldrich: 11836170001).

### Cells lines and tissue culture

The BLaER1 cells and THP-1 cells were maintained in complete RPMI (RPMI containing 10% heat-inactivated fetal calf serum, 1% Pen/Strep and 1% GlutaMAX, all obtained from Thermo Fisher). To trans-differentiate the BLaER1 cells to a macrophage-like cells they were resuspended at 1×10^6^ cells/ml in complete RPMI with 10 ng/ml IL-3, 25 U/ml M-CSF and 100 nM β-Estradiol, and then 100ul was plated in poly-L-lysine (Sigma, P8920) coated 96 well plates and incubated for 6 days to differentiate the cells. On day 6 the cells were used for experiments. The BLaER1 CRISPR KO cell lines were obtained from the laboratory of Prof. Veit Hornung and were generated as described in [23]. THP-1 cells were differentiated in full medium containing 50 ng/mL PMA. The THP-1 CRISPR KO cells were generated using transduction with lentiviruses based on pLenti CRISPR v2 [38] using sgRNAs: Caspase-1-TACCATGAGACATGAACACC, ASC – GCTGGATGCTCTGTACGGGA, NLRP3 - CAATCTGAAGAAGCTCTGGT and Pyrin – TCTGCTGGTCACCTACTATG, followed by selection in 0.75 µg/mL puromycin. Monoclonal cell lines were generated by limited dilution and verified by Sanger sequencing and immunblot.

### Primary cell isolation and differentiation

Monocytes were purified from buffy coat preparations from healthy donors (obtained according to protocols accepted by the institutional review board at the University Clinic Bonn; local ethics votes Lfd. Nr. 075/14). All donors were anonymous. The buffy coat was mixed in a 2:3 ratio with PBS, layered onto a ficoll gradient, and centrifuged at 700g for 20 min without brake. The PBMC layer was extracted from the interface. After washing, it was incubated with CD14 conjugated magnetic beads (Milltenyi Bioscience) and purified using MACS columns (Milltenyi Bioscience) as per the manufacture’s protocol. The cells were then counted and resuspended for direct use or cultured for 3 days in RPMI containing 50 U/ml M-CSF at 2×10^6^ cells/ml to generated hMDM. After the three-day differentiation, hMDM were harvested and plated for experiments, then left to adhere overnight in RPMI containing 25 U/ml M-CSF.

### Inflammasome stimulation assays

Primary monocytes/hMDM: Cells were harvested and seeded the day before the assay. Before the experiment, the media was removed, and fresh media with or without a TLR stimulus was added (LPS 10ng/ml, Pam3CS4K 25 ng/ml) and incubated for 3h. Next, compounds were added and incubated for 15 min (CP-456,773 2.5 uM, VX-765 40 uM, Colchicine 2.5 uM). Inflammasome activators were subsequently added and incubated for 2.5h (TcdB 20 ng/ml, Nigericin 8 µM, PrgI/PA 25 ng/ml, respectively). Plates were centrifuged at 450g for 5 min, then the supernatant harvested for a cytokine or immunoblot analysis, and the cells lysed in RIPA buffer were applicable.

BLaER1 cells: Differentiated BLaER1 cells were seeded in 96 well plates pre-coated in the poly-L-lysine. Before the experiment, the media was removed and fresh media with or without LPS (100 ng/ml) and incubated for 3h. Next, compounds were added and incubated for 15 min (CP-456,773 2.5 µM, VX-765 40 µM, Colchicine 2.5 µM). Inflammasome activators were subsequently added and incubated for 2.5h (TcdB 20 ng/ml, Nigericin 8 uM, PrgI/PA 25 ng/ml, respectively). Plates were centrifuged at 450g for 5mins, and then the supernatant was harvested for cytokine analysis.

THP1s were seeded in 24 well plates in the presence of 50 ng/mL PMA (2·10^5^ per well). Medium was replaced after 16 h; 24 h after this, cells were primed with 100 ng/mL LPS for 3h, followed by treatment with 8 uM Nigericin (NLRP3) or 100 ng/mL PA + 200 ng/mL LFn-PrgI (needletox, NLRC4) for 1.5 h, or 2 µg/mL TcdB for 8h. Supernatants were cleared by centrifugation at 4° C, 1000g for 10 min and IL-1β and TNFα levels were quantified by ELISA. Where indicated, cells were treated with 2.5 µM CP-456,773 (CRID3, MCC950), or 40 µM Vx-765 30 min before and during stimulation.

### Sample preparation and immunoblotting

The supernatant from primary human monocytes, hMDM, or BMDMs (2×10^6^ cells/well in a 6 well plate) was harvested following inflammasome stimulation and the cells lysed in RIPA buffer (20 mM Tris-HCl, pH 7.4, 150 mM NaCl, 1 mM EDTA, 1% Triton X-100, 0.1% SDS, 0.5% deoxycholate, cOmplete protease and PhosSTOP (Roche) inhibitor). First, DNA was disrupted by sonication, then the lysate equivalent to 2×10^5^ cells was mixed at a 1:4 ratio with 4x LDS buffer containing 10% sample reducing agent (Invitrogen). Samples were heated at 95°C for 5 minutes and collected by centrifugation before loading.

Protein from supernatants was precipitated by adding an equal volume of methanol and 0.25 volumes of chloroform, centrifuged for 3 min at 13,000g. Next, the upper phase was discarded, the same volume of methanol of the previous step was added, and the sample was centrifuged for 3 min at 13,000g. The pellet was then dried and resuspended in 1x LDS-sample buffer containing a 10% sample reducing agent (Invitrogen). Samples were heated at 95°C for 5 minutes and collected by centrifugation before loading.

Proteins were separated by 4–12% SDS-PAGE in precast gels (Novex; Invitrogen) with MOPS buffer for proteins above 50kDa or MES buffer for proteins below 50kDa (Novex; Invitrogen). Proteins were transferred onto Immobilon-FL PVDF membranes (Millipore), and nonspecific binding was blocked with 3% BSA in Tris-buffered saline for 1 h, followed by overnight incubation with specific primary antibodies in 3% BSA in Tris-buffered saline with 0.1% Tween-20. For the phosphor-Pyrin immunoblots the transferred membranes were instead blocked in Tris-buffered saline containing 1% milk powder.

Primary antibodies were used as follows: NLRP3 (1:5,000; Cryo-2), human caspase-1 for lysate analysis (1:1,000; Bally-1), murine caspase-1 (1:1000; casper-1) from Adipogen, Pyrin (1:1000, MEFV polyclonal 24280-1-AP) from Proteintech, phospho-Pyrin S241 (1:500; ab200420) from Abcam, human IL-1β (1:1000, AF-201-NA) and murine IL-1β (1:1000, AF-401-NA) from R&D bioscience, Rac (1:1000, clone 102) from BD transduction laboratories, human caspase-1 for supernatant analysis (1:1000, D57A2), Rac1/2/3 (1:1000, rabbit polyclonal #2465) from CST, actin (mouse or rabbit, both 1:1,000 dilution) from LI-COR Biosciences. Membranes were then washed and incubated with the appropriate secondary antibodies (IRDye 800CW, IRDye 680RD or HRP; 1:25,000 dilution; LI-COR Biosciences). In the case of caspase-1 detection or Pyrin detection, the membranes were incubated with washed and analyzed with an Odyssey CLx imaging system (LI-COR Biosciences) and ImageStudio 3 Software (LI-COR Biosciences). For the phosphor-Pyrin immunoblots the membrane was developed using western lighting plus-ECL and analysed on a VersaDoc (Biorad).

### Cytokine measurements

Cytokines were measured either by ELISA or by HTRF as per the manufactures’ instructions. The kits used for ELISA were the human IL-1β (DY201), human TNFα (DY210), mouse IL-1β (DY401) or mouse TNFa (DY410), all from RnD biosystems. HTRF kits used were human IL-1β (62IL1PEC) or human TNFα(62TNFPEB). All assays were read using a SpectraMAX i3 (molecular devices) using the additional HTRF cartridge. Human IL-18 measurements were performed using a cytokine bead array that was generated in our laboratory. The xMAP antibody coupling kit from Luminex (40-50016) was used to conjugate the capture IL-18 antibody (D0044-3, MBL) to the beads. IL-18 was then measured following the standard Luminex protocol and measured on a Magpix multiplexing unit (Luminex).

### siRNA transfection

hMDM were harvested by centrifugation (350g, 5 min) and washed twice in PBS. The cells were then aliquoted to have 1.2 x 10^6^ cells per reaction and centrifuged again for 2 min at 3000 rpm. The supernatant was discarded, and the cell pellet resuspended in 10.5 μL Buffer R with siRNA at 10nM. 10 μL of reaction mix was loaded into the neon electroporation, and the pipette plugged into place within the electroporation tube containing 3ml Buffer E. The electroporation settings were: 1400 V, 20 ms and 2 pulses. Subsequently, the cells were transferred into 2 mL pre-warmed antibiotic-free RPMI. After counting, the appropriate number of cells was seeded in 12-well or 96-well tissue culture plates and incubated for 24 h before experiments.

### Immunofluorescence and microscopy

Following treatment were washed once in PBS, then fixed in 2% PFA at 4C overnight (for BLaER1 ASC speck analysis) or 4% PFA on ice for 30 min. The cells were then washed twice in PBS containing 20 mM Glycine, then twice in PBS. To stain for intracellular targets, the cells were then permeabilized in 0.1% Triton X-100 for 5 min, then blocked in intracellular staining solution (PBS +10% goat serum, 1% HI-FBS and 0.1% Triton X-100) for 30 mins RT. Next, we used Alexa-647 conjugated Phalloidin (Invitrogen, A22287) for 30 min RT in an intracellular staining solution to stain actin. The cells were then washed (3x 5min) and incubated with DAPI (1μg/ml, 10min) before being washed and imaged. For ASC speck detection, the fixed cells were incubated with DRAQ5 (eBioscience, 65-0880-96) for 5 min (1:2000 dilution), then imaged directly. All imaging was performed with an Observer.Z1 epifluorescence microscope, 20× objective (dry, PlanApochromat, NA 0.8; ZEISS), Axiocam 506 mono, and ZEN Blue software (ZEISS). Image analysis of all ASC speck experiments was done using a cell profiler pipeline optimized to detect either ASC specks or nuclei. A minimum of 6 images was analyzed for each condition in each experiment.

### Retroviral transduction and fluorescent activated cell sorting

To produce the virus-containing supernatant 0.4×10^6^ HEK293T cells were plated in 2 mL complete DMEM in one well of a 6 well dish. After 16 - 24 h, HEK293T cells were transfected with retroviral constructs encoding the gene of interest (2 µg per well), the retroviral packaging plasmids gag-pol (1 µg well), and VSV-G (100 ng/well) using GeneJuice transfection reagent (Novagen, 70967). Cells were incubated at 37C, 5% CO2 for approximately 12 h, and then the media was exchanged with RPMI containing 30% HI-FBS, and cells were incubated for another 34 h. After 36 h, the viral supernatant was collected using a 10 mL Luer-lock syringe attached to a blunt 18G needle and then filtered using a 0.45 mm filter unit into a 50 mL falcon. The medium on target cells was removed, and viral supernatant was added to the cells at a 2:1 ratio with complete RPMI. 8 µg/ml polybrene was then added to the diluted virus-containing supernatant. The cells were then centrifuged at 800g for 45 min at 37°C, then harvested and plated in 24 well plates before being incubated for approximately 24 hr at 37°C, 5% CO2. Following incubation, the cells were collected by centrifugation, and the virus-containing medium was removed and replaced by complete RPMI. Transduced cells were passaged three times before frozen stocks were prepared. Cells were sorted for equal expression of Pyrin, and NLRP3 variants using fluorescence assisted cell sorting on a FACS Aria cell sorter for equivalent expression of the fluorescent protein used as a marker of transduction.

### C.difficile co-culture and supernatant generation

Experiments were performed with *C. difficile* DSM 28645 and DSM 29688 obtained from the German Collection of Microorganisms and Cell Cultures (DSMZ, Braunschweig, Germany). Main cultures were cultivated in RPMI 1640, supplemented 10% FBS, 0.014 mM iron sulfate, 4.16 mM cysteine, 4.33 mM proline, 1.11 mM valine, 1.12 mM leucine, 0.72 mM isoleucine, 0.22 mM tryptophane, 0.57 mM methionine and 0.22 mM histidine at 2% O_2_, 5% CO_2_, 37°C, 40-50% humidity using O2 Control InVitro Glove Box (Coy Labs, USA). PBMCs were isolated from three different donors and differentiated into hMDM as described above. Two days before the experiment, cells were seeded in RPMI medium supplemented with 10% HI-FBS at 3.3×10^5^ cells per well in 24 well plates and incubated at normoxic conditions at 37°C for 24 h. On the following day, the cells were placed into a hypoxia chamber (2% O_2_, 5% CO_2_, 37°C, 40-50% humidity) for another 24 h. On the same day, *C. difficile* main cultures of DSM 28645 (toxin-producing) and DSM 29688 (non-toxigenic) were inoculated at an OD_600nm_ of ∼0.01 and incubated for 24 h. On the day of the experiment the medium was removed, and the cells were washed with 500 µl PBS. Following, the cells were treated with 375 µl RPMI or 375 µl RPMI containing 10 ng/ml LPS for 2 h. The OD_600nm_ of both *C. difficile* cultures was determined, and the number of bacterial density was determined using the following formulation:

### *C. difficile* per ml main culture = 26,445,593 x OD_600nm_

The cells were centrifuged at 2500g for 10 min. After centrifugation, the supernatant was passed through a 0.2 µM sterile filter. The pellet was resuspended in RPMI supplemented with 5 mM butyrate and with lower concentrations of glycine (reduced to 0.033 mM), cysteine (0 mM), proline (0 mM), isoleucine (0.095 mM), leucine (0.095 mM), methionine (0.026 mM), serine (0 mM), threonine (0.042 mM) and valine (0.042 mM) and the bacterial density was adjusted to a multiple of infection (MOI) of 300. The cells were treated with 375 µl sterile-filtered *C. difficile* supernatant, living *C. difficile* (300 MOI) or RPMI. The cells were additionally treated with 10 ng/ml LPS, 2 µM CP-456,773 or a combination of both. A lysis control was included by the addition of 0.5% triton-X100 in RPMI. After 3 or 6 hours, the cell supernatant was collected, centrifuged for 10 min at 2500g and frozen at −80°C. The cells were washed two times with 750 µl PBS and lysed by the addition of 80 µl M-PER with cOmplete Mini Protease Inhibitor for 5 minutes. The lysed cell suspensions were collected and stored at −80°C.

### Caspase-1 activity assay

The caspase-1 activity assay was performed using the Caspase-Glo 1 inflammasome assay from Promega (G9951) as per the manufactures instructions. Briefly, cell-free supernatant from inflammasome stimulated cells was mixed with equal amounts for reconstituted caspase-1 reagent and the Luminescence signal read on a SpectraMAX i3 (molecular devices) at 30, 60 and 90 min post mixing.

### qPCR

qPCR quantifications were performed essentially as previously described [39] with the following changes: 500 ng of RNA was used for the RT-PCR and the qPCR was performed using QuantStudio 6 PCR System (Thermo Fisher Scientific). The primer sequences were as follows: Hprt, forward 5′-TCAGGCAGTATAATCCAAAGATGGT-3′ and reverse 5′-AGTCTGGCTTATATCCAACACTTCG-3′; MEFV, forward 5′-GGAAGGCCACCAGACACGG-3′ and reverse 5′-GTGCCCAGAAACTGCCTCGG-3′.

## Acknowledgements

We thank C.C. Kolbe for setting up and optimising the IL-18 Cytokine Bead Array assay, H. Tatge for purification of the different *C.difficile* toxins as well as both the Microscopy Core Facility and Flow Cytometry Core Facility at the University of Bonn for providing help, service and devices funded by the Deutsche Forschungsgemeinschaft (DFG – German Research Foundation) – Projektnummer 13123509. This work was funded by the Deutsche Forschungsgemeinschaft (DFG, German Research Foundation) – Project-ID 369799452 (TRR237) (E.L.), Project-ID 414786233 (SFB1403) (E.L.), Project-ID 432325352 (SFB1454) (E.L.), DFG (GE1017/5-1) (R.G) and Germany’s Excellence Strategy – ImmunoSensation2 – Project-ID 390873048 (EXC2151) (E.L.), The work was also supported by the Helmholtz-Gemeinschaft, Zukunftsthema ‘Immunology and Inflammation’ (ZT-0027) (E.L.) and the German-Israeli Foundation for Scientific Research and Development, Grant No: 1085 (V.H.) This project was partially funded by the Niedersächsiches Ministerium für Wissenschaft und Kultur (MWK), Niedersächsisches Vorab [grant number ZN3380] (K.H.)

---

**Supp figure 1.**
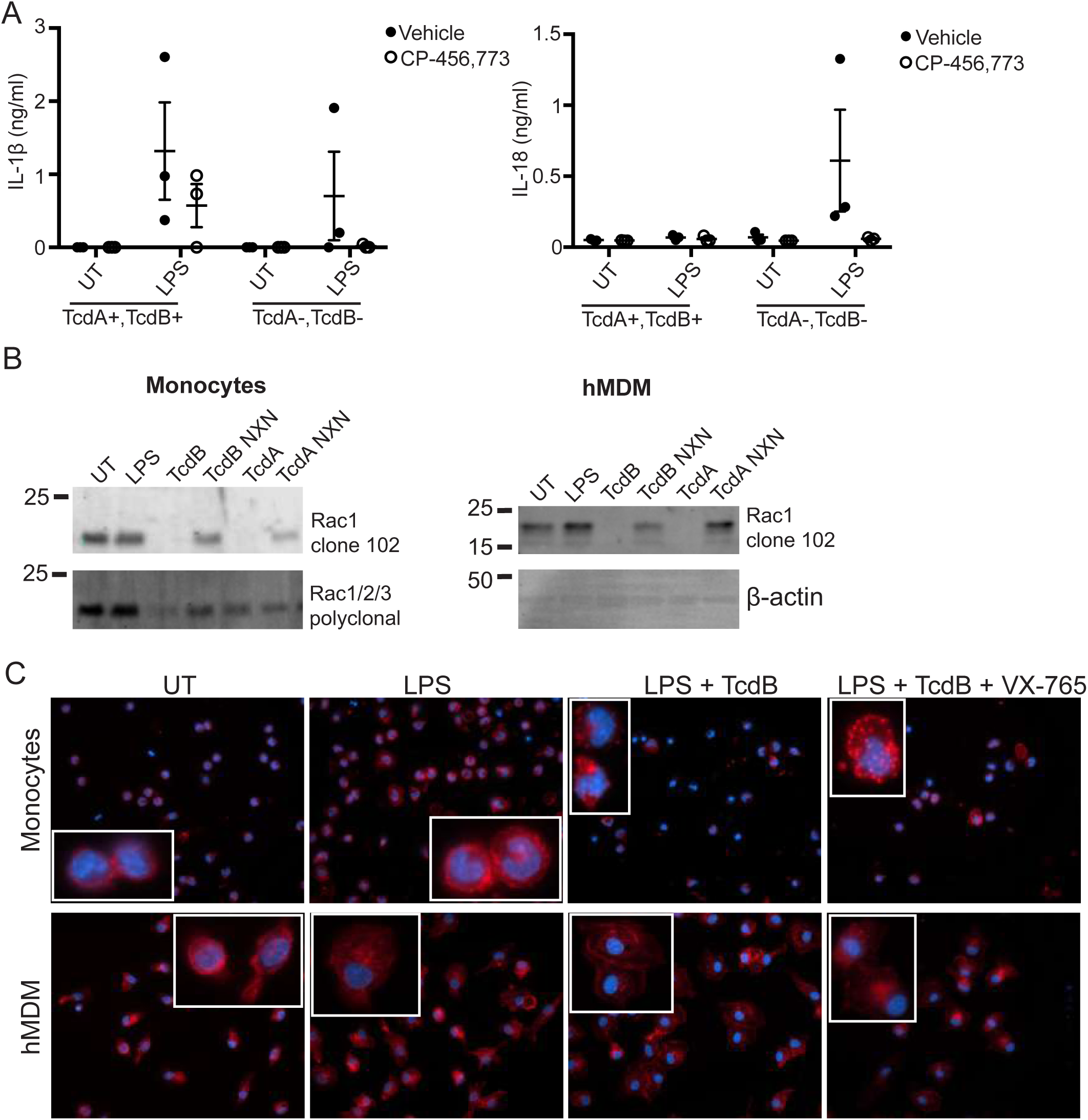
(A) hMDM were incubated with either toxin proficient (TcdA+,TcdB+) or toxin deficient (TcdA-,TcdB-) *C. difficile* (MOI 100:1) for 6h, then the supernatant assessed for IL-1b and IL-18. Mean and SEM of three experimental replicates shown. (B) Rac glucosylation status in either monocytes or hMDM following treatment with the listed toxins (NXN variants - lack glucosyltransferase activity). Representative of 3 experiments. (C) Actin staining following incubation of monocytes or macrophages with or without LPS and TcdB. Treated cells were fixed and stained with Phalloidin 647 to detect actin (red) or with DAPI to detect nuclei (blue). Images are representative from 3 separate donors.

**Supp figure 2.**
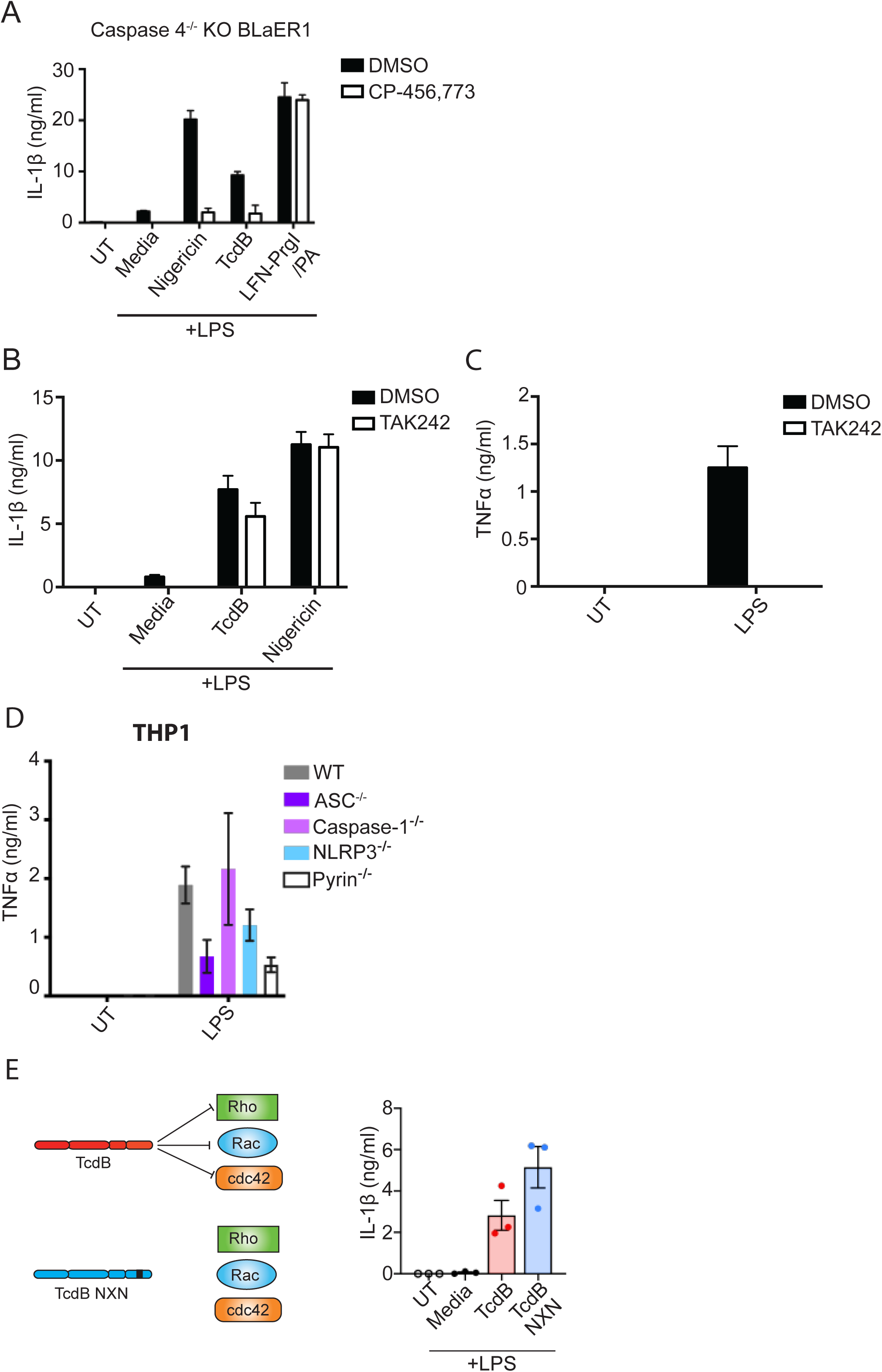
A) Differentiated Casapse-4 deficient BLaER1 cells were stimulated as in Fig 2A. IL-1β was assessed from the harvested supernatants. (B) LPS-primed differentiated WT BLaER1 cells were pre-incubated with TAK242 (2uM, 30 min) then activated with TcdB (20ng/ml) or Nigericin (8uM) for 2h. Harvested supernatant was assessed for IL-1β. (C) Differentiated WT BLaER1 cells were pre-incubated with TAK242 then stimulated with LPS for 4h. TNFa was assessed from the supernatant. (D) TNFa was measured for THP-1 cells from Fig. 2G. Mean and SEM shown for three independent experiments. (E) LPS primed (10ng/ml, 3h) human macrophages were treated either TcdB or the TcdB NXN mutant lacking glucosyltrasferase activity (20ng/ml, 2.5h). Supernatant was harvested and assessed for IL-1β. For A-C the mean and SD of three technical replicates shown, representative of 3 independent experiments. For D and E. the mean and SEM shown for three independent experiments.

